# Phage transcriptional regulator X (PtrX)-mediated augmentation of toxin production and virulence in *Clostridioides difficile* strain R20291

**DOI:** 10.1101/2023.08.24.554564

**Authors:** Jun-Jia Gong, I-Hsiu Huang, Shu-Wei Su, Si-Xuan Xie, Wei-Yong Liu, Cheng-Rung Huang, Yuan-Pin Hung, Shang-Rung Wu, Pei-Jane Tsai, Wen-Chien Ko, Jenn-Wei Chen

## Abstract

*Clostridioides difficile* is a Gram-positive, anaerobic, and spore-forming bacterial member of the human gut microbiome. The primary virulence factors of *C. difficile* are toxin A and toxin B. These toxins damage the cell cytoskeleton and cause various diseases, from diarrhea to severe pseudomembranous colitis. Evidence suggests that bacteriophages can regulate the expression of the pathogenic locus (PaLoc) genes of *C. difficile*. We previously demonstrated that the genome of the *C. difficile* strain RT027 (NCKUH-21) contains a prophage-like DNA sequence, which was found to be markedly similar to that of the φCD38-2 phage. In the present study, we investigated the mechanisms underlying the φNCKUH-21-mediated regulation of the pathogenicity and the PaLoc genes expression in the lysogenized *C. difficile* strain R20291. The carriage of φNCKUH-21 in R20291 cells substantially enhanced toxin production, bacterial motility, biofilm formation, and spore germination in vitro. Subsequent mouse studies revealed that the lysogenized R20291 strain caused a more severe infection than the wild-type strain. We screened three φNCKUH-21 genes encoding DNA-binding proteins to check their effects on PaLoc genes expression. The overexpression of NCKUH-21_03890, annotated as a transcriptional regulator (phage transcriptional regulator X, PtrX), considerably enhanced toxin production, biofilm formation, and bacterial motility of R20291. Transcriptome analysis further confirmed that the overexpression of *ptrX* led to the upregulation of the expression of toxin genes, flagellar genes, and *csrA*. In the *ptrX*-overexpressing R20291 strain, PtrX influenced the expression of flagellar genes and the sigma factor gene *sigD*, possibly through an increased flagellar phase ON configuration ratio.

**Author Summary:** *Clostridioides difficile* is a Gram-positive, spore-forming anaerobic bacterium that can lead to antibiotic-associated diarrhea and pseudomembranous colitis. During the *C. difficile* infection (CDI), the major virulence factor is the secretion of two exotoxins, toxin A and B, to destroy host intestinal epithelium cells. The investigation of bacteriophages affecting the toxicity of *C. difficile* has increasingly been research. We previously isolated a *C. difficile* clinical strain NCKUH-21, which carried a phage-like DNA sequence, and named it φNCKUH-21. However, whether this prophage could enhance the virulence of *C. difficile* and the mechanism for regulating the pathogenicity are still unclear. We successfully created a φNCKUH-21-lysogenized R20291 strain and showed that lysogenized R20291 performed stronger pathogenicity than the wild-type R20291. We found that a φNCKUH-21-specific protein (encoded by *NCKUH-21_03890* gene) might influence *C. difficile* flagellar phase variation to promote toxin production further. These findings are expected to clarify the mechanism for controlling the pathogenicity of φNCKUH-21-infected *C. difficile*. Moreover, we also believe that the existence of hypervirulent *C. difficile* strains carrying a prophage should be monitored proactively in hospitals to prevent severe CDI.

## Introduction

*Clostridioides difficile* is an anaerobic, Gram-positive, spore-forming bacterium that is generally found in the human gut (1, 2). On average, 15,000–30,000 annual deaths in the United States are associated with *C. difficile* infection (CDI); furthermore, CDI-associated annual medical costs exceed US $4.8 billion (1, 2).

The pathogenesis of CDI begins with the bile salt–induced germination of *C. difficile* spores into vegetative cells; in individuals with prolonged antibiotic use, the balance of the endogenous flora in the gastrointestinal tract is disrupted, which allows the newly produced *C. difficile* to gain a growth advantage (3, 4). *C. difficile* has peritrichous flagella, facilitating bacterial attachment onto the host cell surface and mediating swimming motility (5). Furthermore, the biofilm produced by *C. difficile* helps the bacterium effectively colonize the gut epithelium cells of the host (6-9). In *C. difficile*, the expression of genes on the pathogenicity locus (PaLoc) cause the production and secretion of two major exotoxins toxin A (TcdA) and toxin B (TcdB) (10, 11). Some hypervirulent strains of *C. difficile*, such as NAP1/027, produce more significant amounts of toxin to cause more severe CDI symptoms (12). The exotoxins are recognized by specific receptors on intestinal epithelium cells, which induces a downstream signaling pathway, eventually leading to the destruction of the cell cytoskeleton, causing mild to severe chronic diarrhea, pseudomembranous colitis, and even death (13, 14).

Recently, bacterial phase variation has been demonstrated as a critical process regulating bacterial pathogenicity by generating considerable phenotypic heterogeneity (15-17). Upon exposure to selection pressure, phase variation mediated by site-specific DNA recombination can alter the production of bacterial flagella, pili, and exopolysaccharides to evade the host immune response and environmental stress (18-21). The Cdi4 invertible element (flagellar switch) of *C. difficile* regulates the expression of the *flgB* operon through flagellar phase variation (18, 22) to produce the sigma factor SigD. This switch then induces the expression of downstream genes and *tcdR*, which codes for an alternative sigma factor that positively regulates the expression of toxin-related genes (22, 23).

Carbon storage regulator (CsrA) is a small homodimeric RNA-binding protein that its gene locates on *sigD*-dependent operon and serves as a posttranscriptional regulator (24). CsrA influences various processes associated with bacterial pathogenicity, such as flagella synthesis, motility, biofilm formation, toxin production, and carbon metabolism (25-29). The CsrA system has been extensively studied in both Gram-negative and Gram-positive bacteria. In *Escherichia coli*, this regulator typically binds to the 5′ untranslated region of mRNAs to regulate transcript stability (30, 31). Furthermore, CsrA modulates the expression of genes related to glycogenesis and gluconeogenesis (32) and biofilm formation (25). CsrA activates the motility of *E. coli* by increasing the stability of *flhDC* mRNA (33). In *Vibrio cholerae*, CsrA is essential for quorum sensing (38) and effective host colonization and disease establishment (28). In Gram-positive bacteria such as *Bacillus subtilis*, CsrA reduces bacterial motility and flagellin translation in the absence of FliW (34). CsrA plays important roles in different strains of *C. difficile* (27, 35). The results of the experiments involving the overexpression and knockout of *csrA* in the strains 630Δerm and R20291, respectively, indicated that CsrA not only influences metabolism but also increases toxin production, bacterial adhesion, and flagellar defects that inhibit bacterial motility (27, 35).

All characterized bacteriophages that can infect *C. difficile* are temperate phages, and temperate phages can progress to the lytic cycle and lysogenic cycle depending on the physiological status of the infected host (36). The DNA fragments of temperate phages are inserted into the host genome or maintained as a plasmid to form a prophage (37). Prophages could be converted to lytic phages and be released from the host in case of infection with other bacterial strains or environmental stress (38, 39). During the lysogenic cycle, prophages may modify the host phenotype, promote the host’s fitness, upregulate the expression of antibiotic resistance genes, and increase the production of pathogenic factors (40, 41). For example, a lysogenic phage markedly upregulates the expression of the Shiga toxin in *E. coli* and enhances the secretion of the cholera toxin from *V. cholerae* (42). Moreover, some lysogenic *C. difficile* strains exhibit high pathogenicity and increased toxin production (2, 43).

We previously discovered a prophage-like sequence in the genome of *C. difficile* ribotype 027 (RT027) strain NCKUH-21; this prophage was named φNCKUH-21 (43). The genomic sequence of φNCKUH-21 is considerably similar to the φCD38-2 genome with some nucleotide differences (2). Previous studies indicated that φCD38-2 exhibits a host range in which 48% of tested isolates were infected, among which 80% corresponded to the RT027 strains (2). Notably, prophage φCD38-2 exists as a circular plasmid in the lysogenic host. A ParA/Soj-like protein, involving plasmid maintenance, was encoded in the genome of φCD38-2 and φNCKUH-21 (2, 43, 44). Moreover, φCD38-2 can upregulate toxin expression in the host bacterium (2), and influence the phase variation of *C. difficile* surface protein gene (*cwpV*) and the expression of metabolic genes (45). However, the mechanisms involved in the prophage-mediated regulation of the pathogenicity mediated by *C. difficile* remain unknown. Therefore, in this study, we investigated the mechanisms underlying the φNCKUH-21-mediated regulation of the pathogenicity and PaLoc genes expression of lysogenized R20291. We found that the carriage of φNCKUH-21 in the lysogenized R20291 strain upregulated the expression of multiple pathogenicity-related genes significantly, particularly those responsible for toxin production, bacterial motility, and biofilm formation. We were able to demonstrate further that PtrX (NCKUH-21_03890) alone, a putative transcriptional regulator encoded by φNCKUH-21, could regulate flagellar and PaLoc genes via the modulation of flagellar phase variation ON/OFF ratio in R20291.

## Results

### φNCKUH-21 induced by the mitomycin C treatment can infect R20291

Previously, we observed a 99% identity match between the genomic DNA sequences of φNCKUH-21 (Fig 1A, S3 Table) and φCD38-2 (43). The genetic organization between these two phages was 100% identical except for some nucleotide differences. To examine whether φNCKUH-21 phage could be induced from strain NCKUH-21, we performed a prophage induction test by using mitomycin C, a widely used phage-inducing antibiotic (46). The optical density (OD) of NCKUH-21 was measured to confirm successful phage induction; the OD value decreased significantly after 3–5 h of mitomycin C treatment (S1A Fig), suggesting bacterial lysis for phage release. Moreover, the complete virus particle of φNCKUH-21 could be observed using transmission electron microscopy (TEM) with negative staining (Fig 1B). To evaluate whether the induced φNCKUH-21 can infect other strains of *C. difficile*, we used the *C. difficile* strain R20291, a representative of the hypervirulent strain BI/NAP1/027, as the host bacterium. After infecting R20291 cells with φNCKUH-21, several colonies were inoculated for PCR checking. The PCR of the genomic DNA using φNCKUH-21-specific primers gp39-F/gp39-R demonstrated that the successful infection of φNCKUH-21 (S1B Fig). There was no significant growth difference between lysogenized and wild-type (WT) R20291 cultured in BHIS broth (S1C Fig). These findings indicated that the mitomycin C treatment of NCKUH-21 could induce φNCKUH-21 production, and the resulting phage particles could infect R20291 cells.

**Fig 1.**
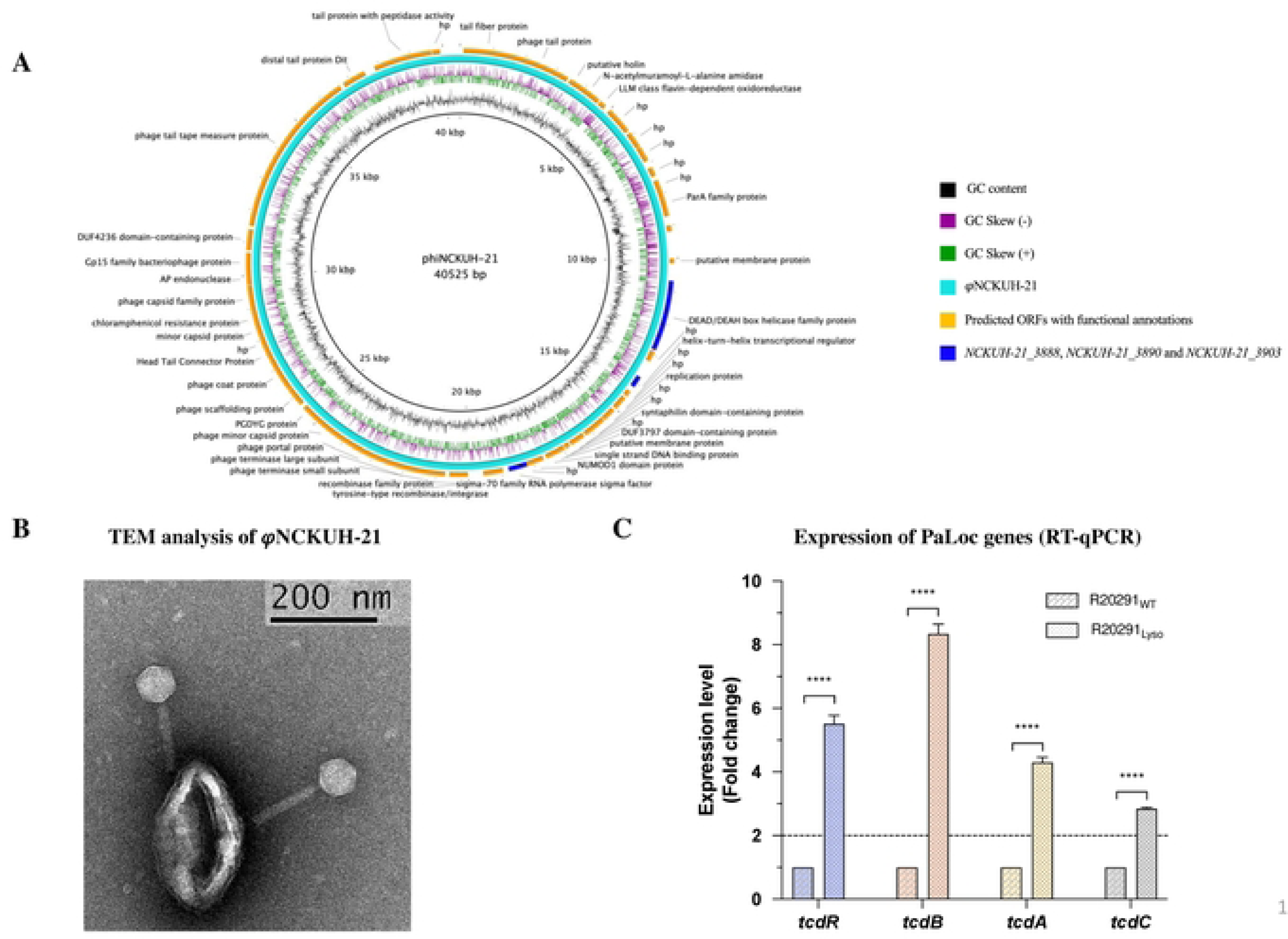
The characteristics of φNCKUH-21 and the PacLoc genes expression in lysogenized R20291. (A) Schematic of the complete φNCKUH-21 genome (40,525 bp) from *C. difficile* strain NCKUH-21 was performed with BRIG. Predicted open reading frames (ORFs) with functional annotations above the respective genes were represented with orange segments. *NCKUH-21_03888*, *NCKUH-21_03890* (*ptrX*), and *NCKUH-21_03903* were represented by blue segments. hp, hypothetical protein. (B) TEM image of negatively stained φNCKUH-21. (C) The relative expression folds of PaLoc genes (*tcdA*, *tcdB*, *tcdR*, and *tcdC*) were determined. Total RNA was extracted from bacterial cultures of R20291_WT_ and R20291_Lyso_ as the OD_600_ value reached 0.8. RT-qPCR and the 2^–ΔΔCt^ method were performed to determine the RNA expression levels. The dotted line indicating the gene expression in the lysogenized strain was two-fold higher than in the WT strain. All data were presented as the mean ± standard deviation values. At least three independent biological replicates of the experiment were conducted with similar results. Data from one representative independent biological replicate are shown. Statistical significance was evaluated using the t-test analysis of variance test (****p ≤ 0.0001).

### Prophage φNCKUH-21 enhances the toxin production and cytotoxicity of lysogenized R20291

Previously, φCD38-2 was shown to induce the production of toxins A and B in lysogenized *C. difficile* bacteria (2). The composition of PaLoc, which contains toxin genes *tcdA*, *tcdB*, and three additional small open reading frames, *tcdR*, *tcdE*, and *tcdC*, is highly conserved in all toxigenic strains of *C. difficil*e (47). To confirm whether the prophage φNCKUH-21 also upregulates the expression of PaLoc genes in the *C. difficile* strain R20291, we performed real-time reverse transcription-quantitative PCR (RT-qPCR) to analyze the relative expression levels of toxin-related genes. As shown in Fig 1C, at OD 0.8 (late log phase), the expression levels of *tcdA* and *tcdB* were 4.1- and 8.2-fold, respectively, higher in lysogenized R20291 than in the WT strain. These findings indicated that φNCKUH-21, like φCD38-2, could upregulate the expression of PaLoc genes in *C. difficile*.

To further investigate whether φNCKUH-21 influences the toxicity of strain R20291, we performed the toxin enzyme-linked immunosorbent assay (ELISA) to quantify the relative amounts of TcdA and TcdB in the supernatants of lysogenized and WT R20291. As shown in Fig 2A, the lysogenized R20291 produced more toxins than the WT strain. To confirm that increased toxin production could have more cellular toxicity, we subjected Caco-2 cells to WT and lysogenized R20291 supernatants, respectively. As shown in Fig 2A, supernatants of the lysogenized R20291 were significantly more toxic than that of the similarly diluted WT supernatant. In addition, the toxicity of the lysogenized R20291 supernatants was non-significantly different from the 2-fold diluted WT strain supernatant when the dilution reached 16-fold. In summary, infection of strain R20291 with the φNCKUH-21 resulted in a notable increase in toxin production and cytotoxicity compared to the WT strain.

**Fig 2.**
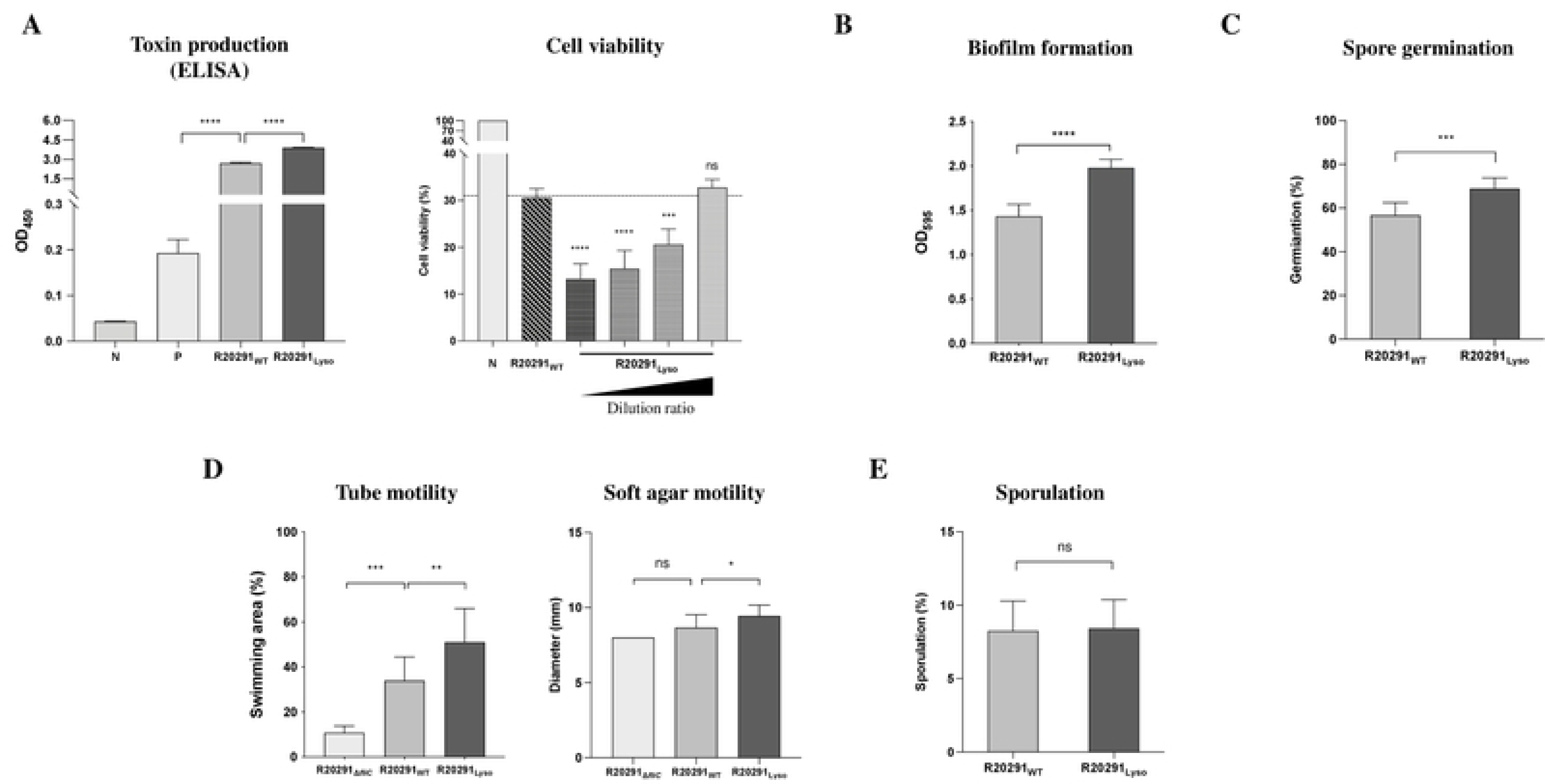
φNCKUH-21 enhances the toxicity and pathogenicity of lysogenized R20291. (A) The harvested bacterial supernatants were 5-fold diluted to perform Toxins ELISA assay for the evaluation of the total amounts of toxins A and B secreted by lysogenized and wild-type R20291 grown in TY broth for 30 hr. N (negative control: sample diluent); P (positive control: inactivated toxin A and B). Bacterial cytotoxicity was assessed using Caco-2 cells (3x10^4^) to coculture with bacterial condition medium, and trypan blue was utilized to calculate the living cells. N (negative control), cell and bacterial culturing media (TY broth) treated groups; R20291_WT_, 2-fold diluted wild-type R20291 supernatant; R20291_Lyso_, serially 2-fold diluted lysogenized R20291 supernatant. Furthermore, the pathogenicity of both R20291_WT_ and R20291_Lyso_ was determined with bacterial (B) spore germination, (C) biofilm formation, (D) swimming motility, and (E) sporulation assays. All data were presented as the mean ± standard deviation values. At least three independent biological replicates of toxin production-associated experiments were conducted with similar results. Data from one representative independent biological replicate are shown. Other pathogenic phenotype results representative of three independent experiments and each performed in triplicate. Statistical significance was evaluated using the t-test and one-way analysis of variance test (\**p* ≤ 0.05; \*\**p* ≤ 0.01; \*\*\**p* ≤ 0.001; \*\*\*\**p* ≤ 0.0001; ns, not significant).

### Prophage φNCKUH-21 enhances the pathogenicity of lysogenized R20291

To understand more about the effects of φNCKUH-21 on the pathogenicity of strain R20291, we next determined whether φNCKUH-21 regulates other pathogenic phenotypes of strain R20291. Biofilm formation is an important factor in the persistence and pathogenesis of CDI (48); therefore, we compared lysogenized and WT R20291 in terms of their biofilm formation ability. After 5 days of growth in BHIS broth containing 0.1 M glucose, lysogenized R20291 exhibited higher biofilm formation ability than WT strain R20291 (Fig 2B). This finding suggested that φNCKUH-21 enhanced the biofilm formation of lysogenized R20291.

We also assessed other pathogenicity-related characteristics, such as effective spore germination, bacterial motility, and sporulation, in both WT and lysogenized R20291 strains. Although there was no significant difference in sporulation (Fig 2E), the spore germination rate was significantly higher in lysogenized R20291 than in the WT strain (Fig 2C). Finally, we evaluated the swimming motility of the two strains. Swimming motility was significantly higher in lysogenized R20291 than in WT R20291 (Fig 2D). In summary, φNCKUH-21 could also enhance biofilm formation, spore germination, and bacterial motility in the lysogenized R20291 strain.

### The lysogenized R20291 strain causes more severe infection in the CDI mouse model than the WT strain

To investigate whether the virulence of lysogenized R20291 is enhanced in the CDI mouse model, we infected C57BL/6 mice with lysogenized R20291 or WT R20291. Mice were administered antibiotics through drinking water for five days and then challenged with *C. difficile* vegetative cells or phosphate-buffered saline (PBS) buffer only in the mock group; mice were euthanized three days after infection (Fig 3A). The decreasing rate of changes in the mouse body weight from day 0 to day 3 post infection, was significantly higher in mice challenged with lysogenized R20291 than in those challenged with WT R20291 or those without any infection (Fig 3B). The survival of mice was shown in Fig 3C, and the mock group exhibited 100% survival. two other groups that were challenged with *C. difficile* vegetative cells. These two groups, which were challenged with the WT and lysogenized R20291 vegetative cells, showed no significant difference in mice survival rate.

**Fig 3.**
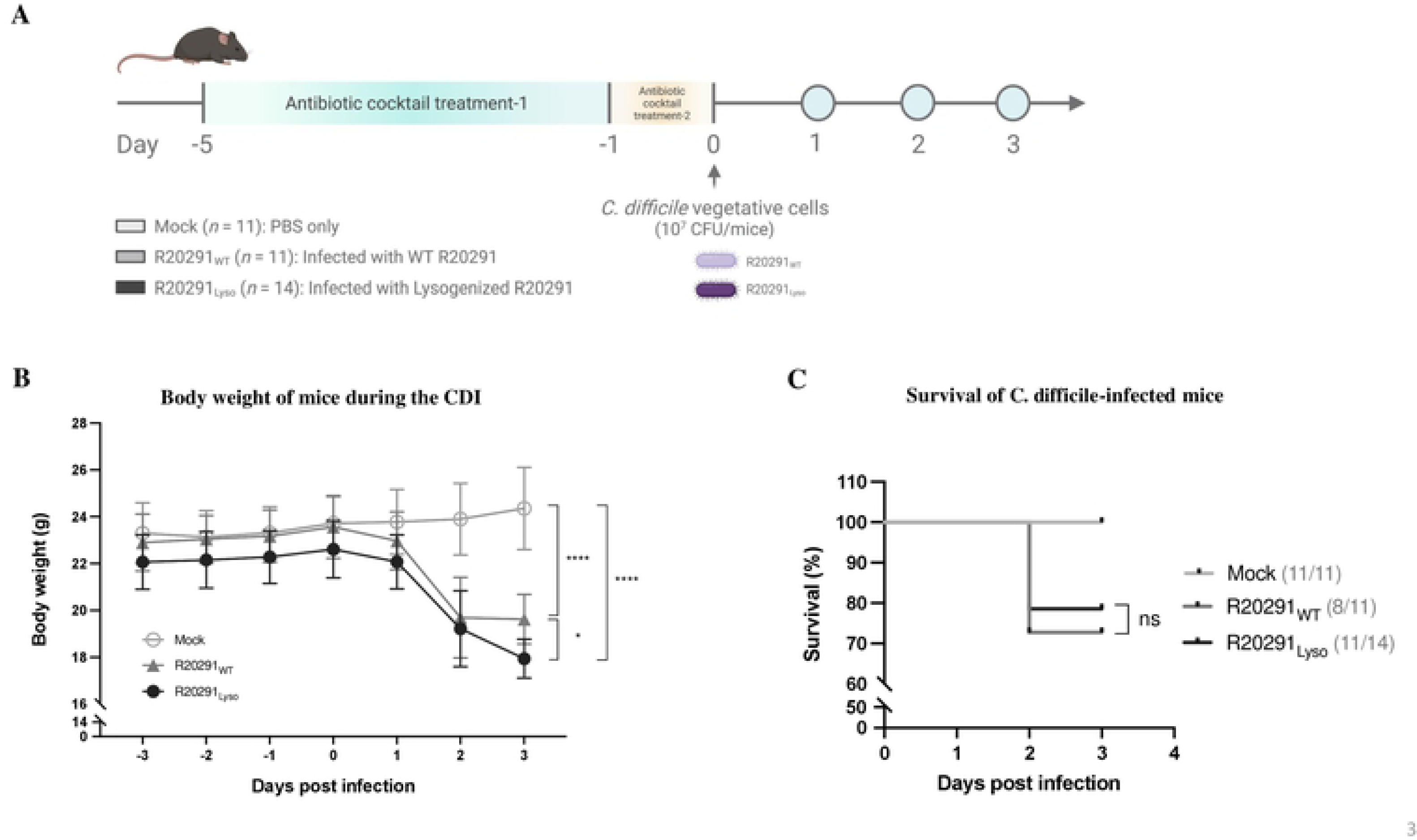
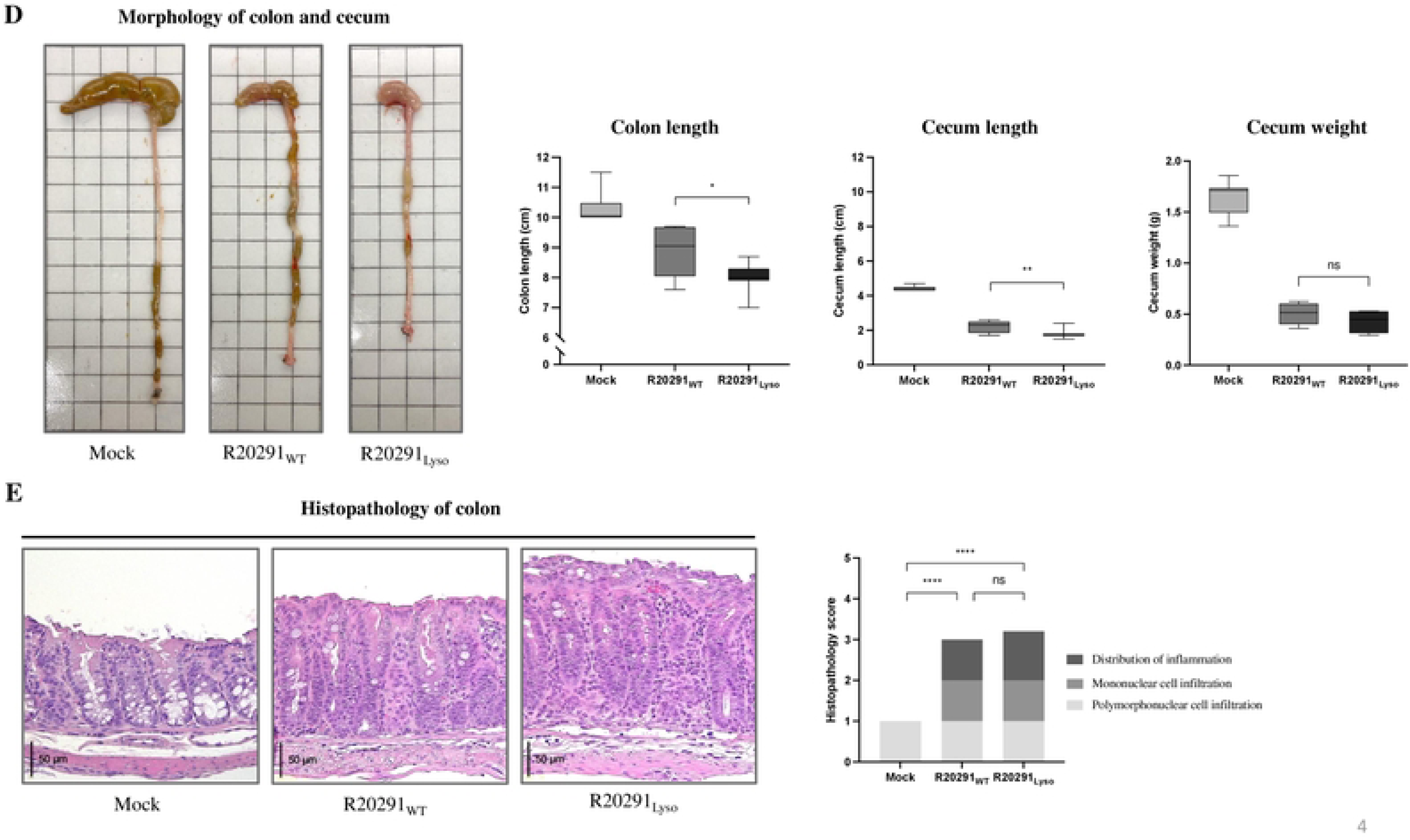
The comparison between wild-type and lysogenized R20291 in mice infection model. (A) Progression schematic of antibiotic-induced *C. difficile* infected mice model (Created with BioRender.com). After five days of antibiotic cocktail treatment, mice were challenged with 10_7_ *C. difficile* vegetative cells and sacrificed at day three post-infection. Mock, mice administered orally with only phosphate-buffered saline (PBS); R20291_WT_, mice administered orally with WT R20291 in PBS; R20291_Lyso_, mice administered orally with lysogenized R20291 in PBS; Antibiotic cocktail treatment-1: containing kanamycin, gentamycin, colistin, metronidazole, and vancomycin; Antibiotic cocktail treatment-2: kanamycin, gentamycin, and colistin. (B) Body weights of mice were monitored during the CDI, and (C) the survival rates of different *C. difficile* strains-challenged mice; (D) colonic tissue morphology, colon length, cecum length, and cecum weight; (E) colonic histopathology and scoring were assessed. All data were presented as the mean ± standard deviation values. Animal experiments were representative of two independent experiments. Symbols represent data of individual mice. Statistical significance was evaluated using the t-test, survival-test, and one-way analysis of variance test (∗*p* ≤ 0.05; ∗∗*p* ≤ 0.01; \*\*\*\**p* ≤ 0.0001; ns, not significant).

The length and weight of the mouse colon were measured to determine the severity of CDI-induced inflammation. Although there was no significant difference in cecal content weight after infecting with both *C. difficile* strains (Fig 3D), mice infected with lysogenized R20291 exhibited a significantly shorter cecum and colon length than did the healthy mice or those challenged with WT R20291 (Fig 3D). The analysis of the intestinal morphology revealed that the infected mice retained less fecal content in the colon lumen than did the uninfected mice (Fig 3D). Histopathological analysis of the colonic indicated that infected mice had more tissue inflammation and mononuclear infiltrated into their colon villus than in uninfected mice (Fig 3E). However, no significant difference was noted in colonic histopathological score between mice infected with WT R20291 and those infected with lysogenized R20291 (Fig 3E). These findings indicated that although the lysogenized R20291 strain exhibited a higher virulence than the WT strain, this increased virulence didn’t impact mouse survival significantly.

### *NCKUH-21_03890* significantly upregulates the expression of PaLoc genes and the pathogenicity of R20291

We next tried to identify which φNCKUH-21 genes were involved in the upregulation of PaLoc genes expression. (43). Based on bioinformatic analysis, three φNCKUH-21 genes, *NCKUH-21_03888*, *NCKUH-21_03890* and *NCKUH-21_03903*, appeared to encode DNA or RNA binding proteins. We hypothesized that phage proteins could regulate the expression of PaLoc genes directly, therefore, these three genes were first used to test. In addition, the DNA sequences of these three genes are 100% identical between φNCKUH-21 and φCD38-2. These three genes were cloned with the shuttle cloning vector pMTL-84151, respectively. Through conjugation assay, the plasmids were transferred from competent *E. coli* cells to the *C. difficile* strain R20291. Bacterial growth curve analysis revealed no significant difference between the transconjugants (Fig 4A). The RT-qPCR analysis results showed there was no more than a two-fold significant difference in PaLoc genes expression between R20291_p3888_ and R20291_pVec_ transconjugants (Fig 4B). The *tcdR* and *tcdA* expression levels in R20291_p3903_ transconjugant were more than two-fold higher than the vector control. Notably, the expression levels of *tcdR*, *tcdB*, *tcdA*, and *tcdC* in R20291_p3890_ were 32.2-, 5.1-, 3.6-, and 29.0-fold, respectively, higher than in the vector control (Fig 4B). Because R20291_p3890_ transconjugant expressed a higher expression level of PaLoc genes than R20291_pVec_, we believe that the protein encoded by *NCKUH-21_03890* plays a significant role in the upregulation of PaLoc genes expression in lysogenized R20291.

**Fig 4.**
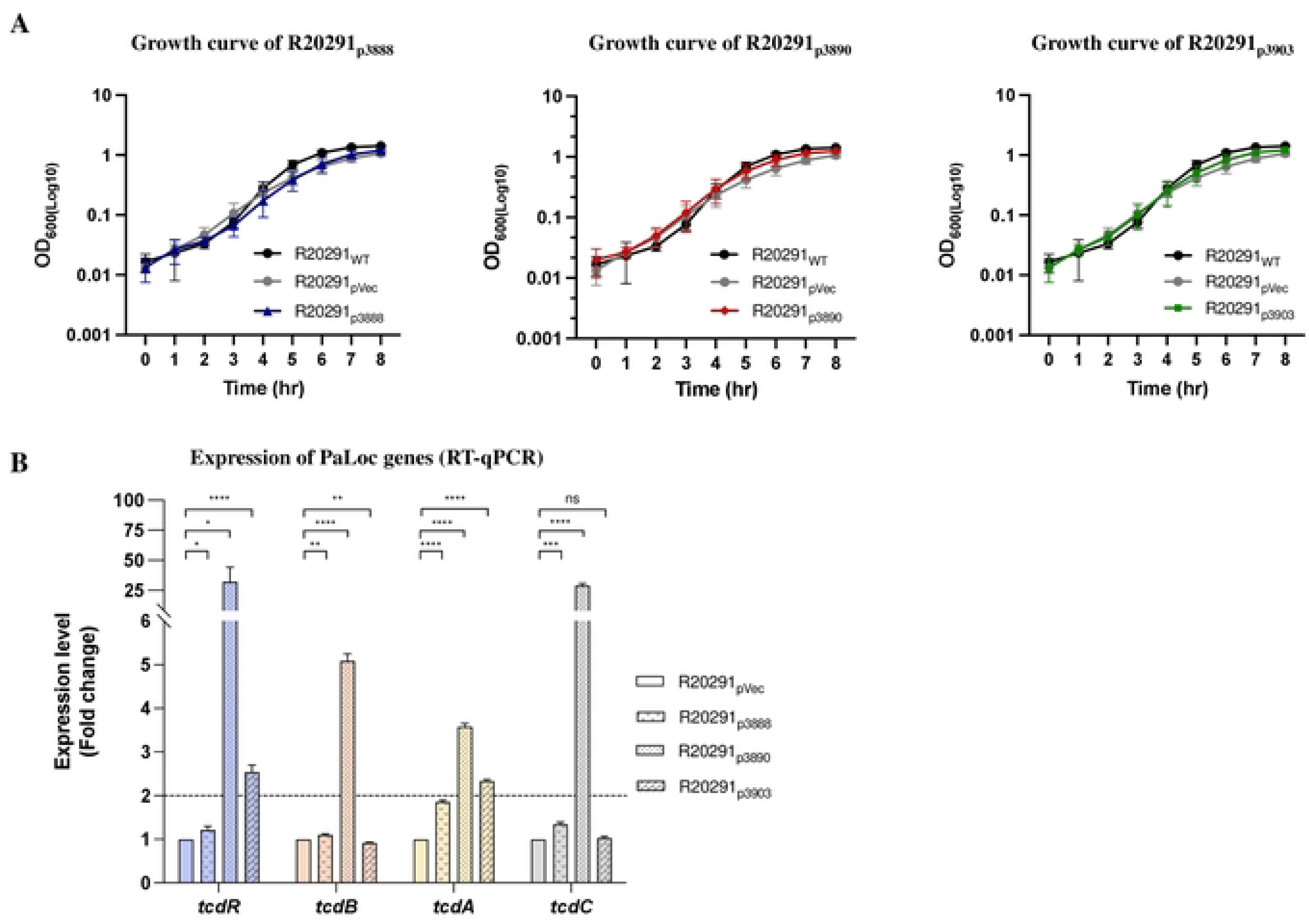
The growth curve and relative expression of PaLoc genes in three phage-gene-overexpressing R20291 strains. (A) The growth curves of three R20291 transconjugants carrying φNCKUH21-specific genes compared with the R20291_WT_ and the vector (pMTL-84151) control. (B) The comparison of PaLoc genes expression between R20291_pVec_ and R20291_p3888_, R20291_p3890_, and R20291_p3903_ transconjugants. The dotted line indicating the gene expression of the R20291 transconjugant was two-fold higher than the R20291_pVec_. All data were presented as the mean ± standard deviation values. At least three independent biological replicates of the experiment were conducted with similar results. Data from one representative independent biological replicate are shown. Statistical significance was evaluated using the t-test analysis of variance test (\**p* ≤ 0.05; \*\**p* ≤ 0.01; \*\*\**p* ≤ 0.001; \*\*\*\**p* ≤ 0.0001; ns, not significant).

To determine the effects of *NCKUH-21_03890* on strain R20291 pathogenic phenotypes, we also compared the biofilm formation, spore germination, and motility of R20291_p3890_ and R20291_pVec_. As shown as the biofilm formation assay was performed to compare the difference between R20291_p3890_ and R20291_pVec_ (Fig 5A), the biofilm formation in R20291_p3890_ transconjugant was significantly enhanced than the vector control. In the spore germination assay, there was no any significant difference between R20291_p3890_ and R20291_pVec_ (Fig 5B). The swimming motility of R20291_p3890_ and R20291_pVec_ was determined using two different methods: tube motility assay and soft agar motility assay. The motility in R20291_p3890_ transconjugant was observed to exhibit significantly higher swimming motility than the vector control (Fig 5C). In summary, our findings suggest that *NCKUH-21_03890* enhances the expression of PaLoc genes, and simultaneously increased the biofilm formation and bacterial motility of R20291. Since *NCKUH-21_03890* was annotated as a helix-turn-helix transcription factor (S3 Fig) and its function is still unclear, we renamed it to <u>p</u>hage transcription regulator X, *ptrX*, and R20291_p3890_ was renamed to R20291_pPtrX_.

**Fig 5.**
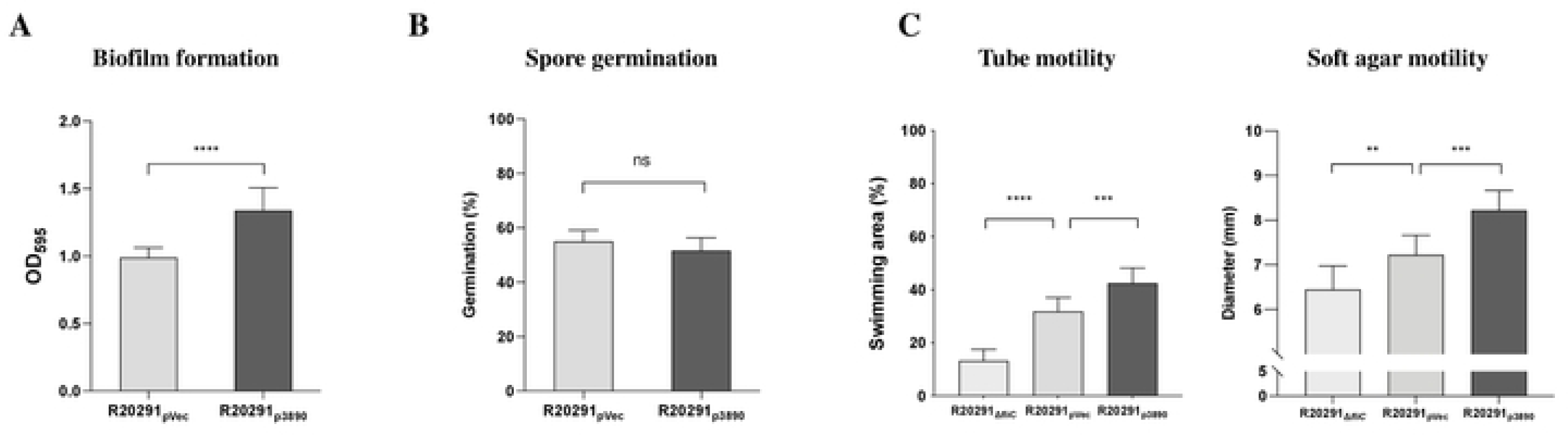
Pathogenic phenotypes of *NCKUH-21_03890*-overexpressing R20291 transconjugant. (A) Heat-treated spores were inoculated in BHIS agar containing 0.1% taurocholate to calculate the spore germination rate. (B) Levels of biofilm formation activity were measured in terms of OD_595_ values. (C) Tube motility and soft agar assays were used to assess bacterial flagellar motility. R20291_ΔfliC_, *C. difficile* R20291 *fliC* mutant strain. All data were presented as the mean ± standard deviation values. The data were representative of three independent experiments, and each performed in triplicate. All statistical analyses were performed using the t-test and one-way analysis of variance test (\*\**p* ≤ 0.01; \*\*\**p* ≤ 0.001; \*\*\*\**p* ≤ 0.0001; ns, not significant).

### Overexpression of PtrX significantly upregulates the expression of the flagellar genes of R20291

To investigate how many *C. difficile* genes are regulated by the DNA-binding protein PtrX and what their function are, we performed a gene expression array analysis, and three independent RNA samples each from R20291_pPtrX_ and R20291_pVec_ were used in the comparison. The differentially expressed genes (DEGs) in this analysis were defined as the gene expression difference more than two-fold between conditions. As shown in Fig 6A, sixty-four DEGs were upregulated in R20291_pPtrX_ transconjugant and twenty-three DEGs were upregulated in the vector control. The bacterial RNA was isolated when the bacterial culture was grown until OD_600_ reached 0.4 to 0.6 (middle log phase) in this study, and the expression of toxin genes responds to nutrient limitation. Therefore, DEGs in this array analysis didn’t include toxin genes is reasonable. The DEGs list was uploaded to STRING for performing protein-protein interaction network functional enrichment analysis, and the gene ontology results in the biological process demonstrated that many DEGs were involved in flagellum assembly, flagellum-dependent cell motility, and chemotaxis (Fig 6B, S4 Table).

**Fig 6.**
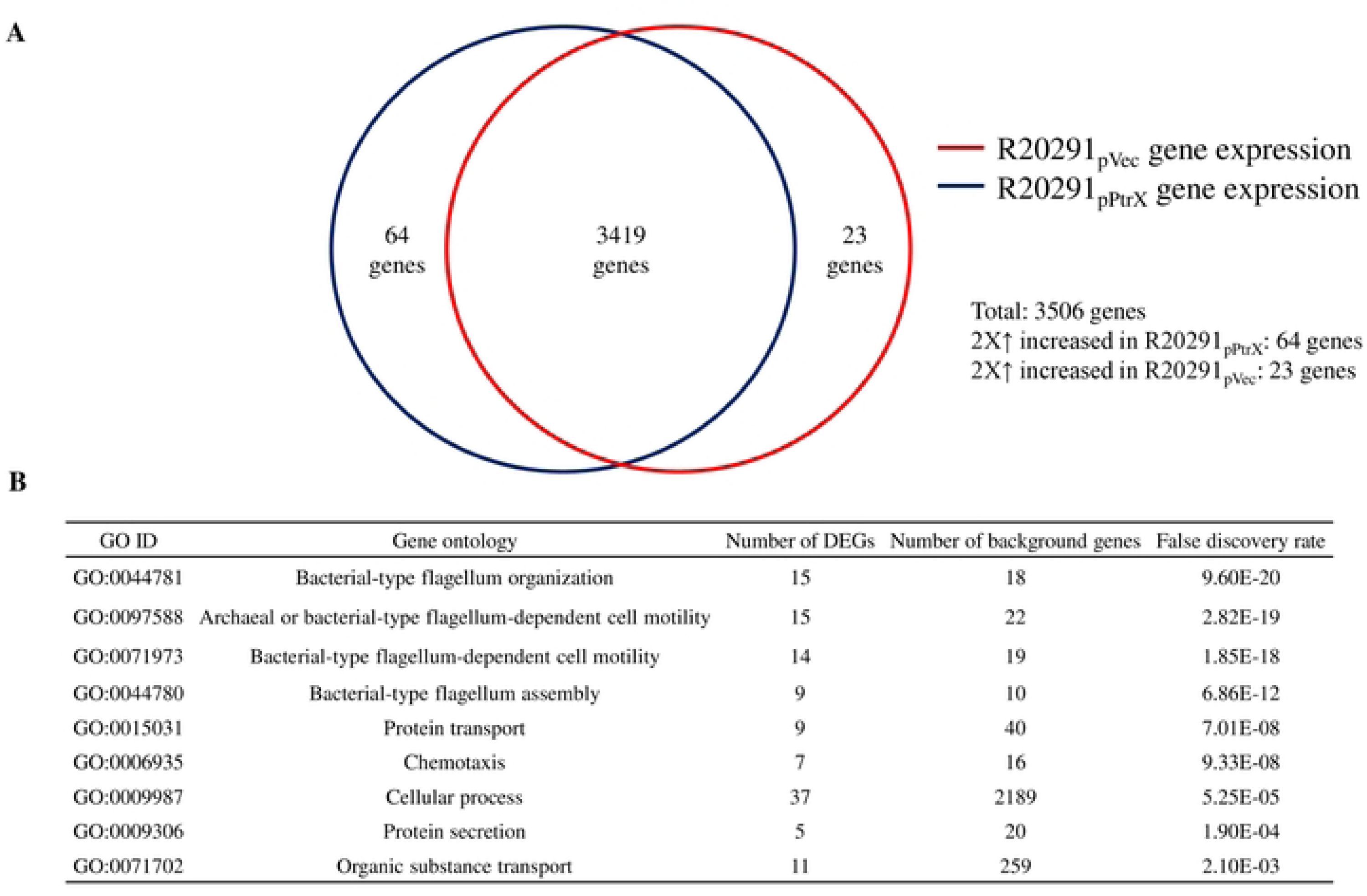
PtrX extensively upregulates *C. difficile* flagellar-related genes in microarray and gene ontology (GO) enrichment analyses. (A) Bacterial gene expression microarray. The p-value adjusted with the false discovery rate (FDR) was used to determine the differentially expressed genes (DEGs). The dark blue circle was the genes expressing in R20291_pPtrX_ transconjugant, 64 genes were 2-fold higher than the R20291_pVec_ transconjugant (the red circle), and 23 genes were 2-fold lower than the vector control. This experiment was performed in three biological replications. (B) The gene ontology (GO) classification in the biological process of DEGs was analyzed by STRING protein-protein interaction networks functional enrichment analysis.

Next, RT-qPCR was performed to verify the results of gene expression microarray analysis. Early-stage flagellar genes (*flgB* operon) are the first to be transcribed. These genes encode the proteins required for the basal body, motor, and rod of the flagella. Although *flgB* was not one of DEGs, many DEGs were encoded by the *flgB* operon. The alternative sigma factor, SigD, is also encoded by the *flgB* operon. SigD not only induces the expression of late-stage flagellar genes but also positively regulates the expression of *tcdR*, which encodes an alternative sigma factor that activates the transcription of *tcdA* and *tcdB* (49). Therefore, the regulation of flagellar genes could influence pathogenicity by modulating the expression of toxin-related genes (S2 Fig). The gene expression microarray analysis (Fig 7A) revealed that the expression level of *tcdA* was 1.48-fold higher in R20291_pPtrX_ transconjugant than in the vector control; moreover, the expression levels of *sigD* and *flgB* were 6.50- and 6.64-fold, respectively, higher than in the vector control. The expression level of *CDR20291_3128* (Cdi6; *C. difficile* inversion site 6), which has previously been indicated that the phase ON configuration performed in the inverted orientation (17), was 2-fold lower in R20291_pPtrX_ transconjugant than in the vector control.

**Fig 7.**
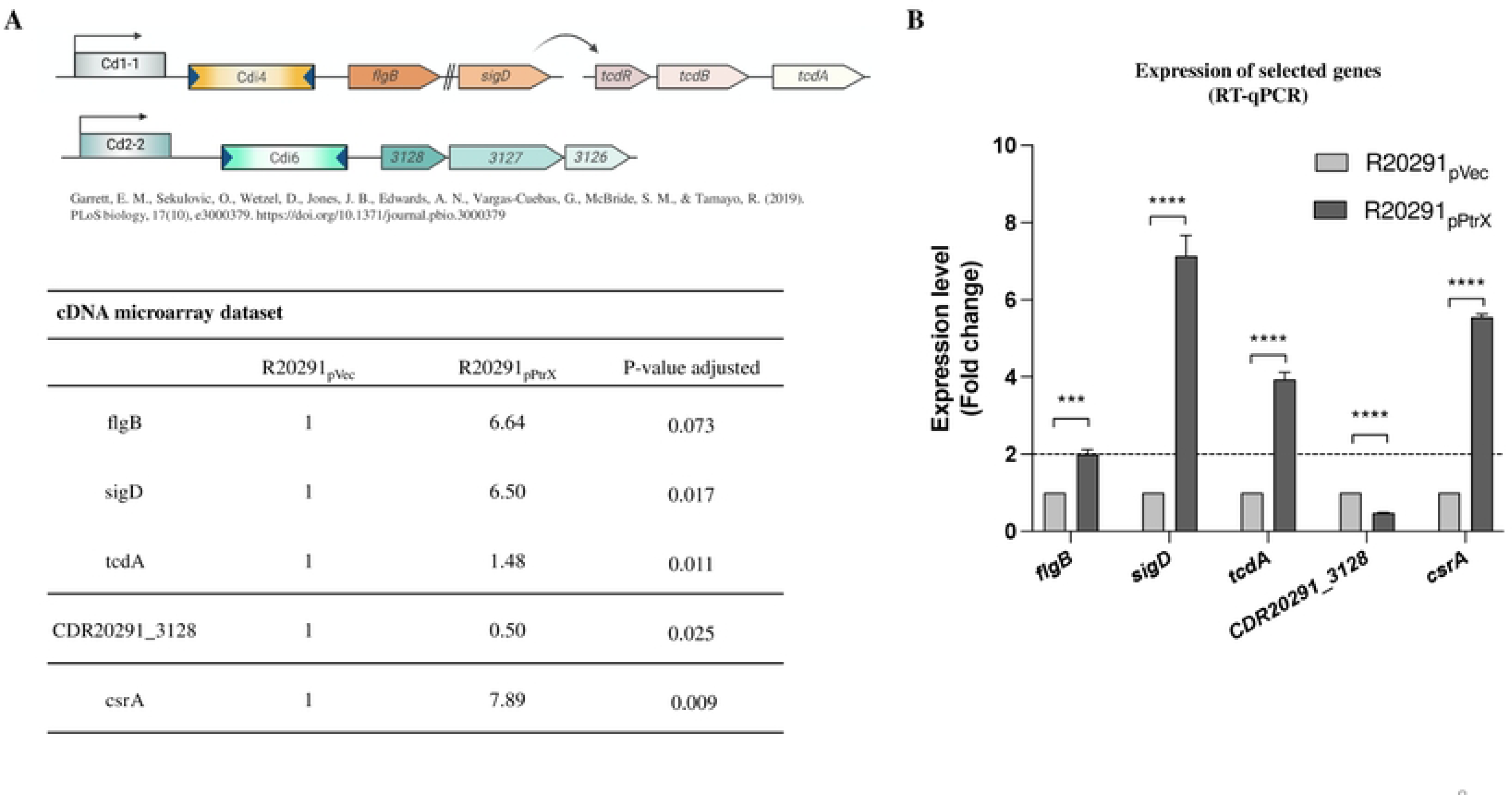
The validation of DEGs in gene expression microarray using RT-qPCR. (A) Schematic diagrams of Cdi4 and Cdi6 invertible DNA elements in *C. difficile* R20291(Created with BioRender.com). Those genes comprised the upregulated genes in the gene expression microarray data of R20291_pPtrX_ transconjugant. (B) RT-qPCR was performed to determine the expression of flagellar-(*flgB*, *sigD*) and virulence-associated (*tcdA*, *CDR20291_3128*, and *csrA*) genes of R20291_pPtrX_ transconjugant. The dotted line indicating the gene expression in the R20291_pPtrX_ group was two-fold higher than in the vector control. All data were presented as the mean ± standard deviation values. At least three independent biological replicates of the experiment were conducted with similar results. Data from one representative independent biological replicate are shown. All statistical analyses were performed using the t-test analysis of variance test (\*\*\**p* ≤ 0.001; \*\*\*\**p* ≤ 0.0001).

As the RT-qPCR results shown in Fig 7B, the expression levels of *flgB*, *sigD*, and *tcdA* in R20291_pPtrX_ were 2.00-, 7.14-, and 3.94-fold higher than in the vector control. By contrast, the expression level of Cdi6 was at least 2-fold lower in R20291_pPtrX_ transconjugant than the vector control. This result provides supportive evidence for the gene expression microarray, and the gene expression of DEGs exhibits a similar trend in two different assays.

### *ptrX* modulates *csrA* expression and flagellar phase variation in R20291

Gene expression microarray analysis revealed that the expression level of another important gene, *csrA*, which is associated with increased *C. difficile* toxin generation and pathogenicity (27), was significantly higher in R20291_pPtrX_ than in the vector control R20291_pVec_ (Fig 7A). In the RT-qPCR analysis, the expression level of *csrA* in the transconjugant was approximately 5.5-fold higher than in the vector control (Fig 7B). Furthermore, the expression level of *csrA* was approximately 2-fold higher in lysogenized R20291 than in WT R20291(S4A Fig).

Based on our gene expression microarray analysis results, *CDR20291_3128* and flagellar genes were included in the DEGs list. Two Cdi sites, Cdi6 and Cdi4, have been reported to regulate the expression of *CDR20291_3128* and *flgB* operon genes during phase variation (50). Therefore, we hypothesized that PtrX might regulate toxins and flagella production and activity via the Cdi4 switch during phase variation. No significant difference was observed in *cwpV* expression between R20291_pPtrX_ transconjugant and the vector control (S4B, C Fig).

To confirm whether the expression of PtrX in R20291 influences the flagellar switch for modulating the expression of PaLoc genes, we extracted genomic DNA from R20291_pVec_, R20291_pPtrX_ transconjugants, WT R20291 and lysogenized R20291. Two previously reported primers, qPCR_FlgSwit-ON, qPCR_FlgSwit-OFF and qPCR_FlgSwit-REV (18), were used to perform the absolute quantification of the Cdi4 phase ON and phase OFF configurations. The percentage of phase ON configuration was approximately 1.7-fold higher in the R20291_pPtrX_ transconjugant (46.4%) than in the R20291_pVec_ (27.1%) (Fig 8A). Furthermore, the percentage of phase ON configuration in lysogenized R20291 (35.8%) was approximately 2.7-fold higher than in WT R20291 (13.3%) (Fig 8B). In summary, the expression of *csrA* and the percentage of Cdi4 phase ON configuration were increased in both lysogenized R20291 and R20291_pPtrX_ transconjugant.

**Fig 8.**
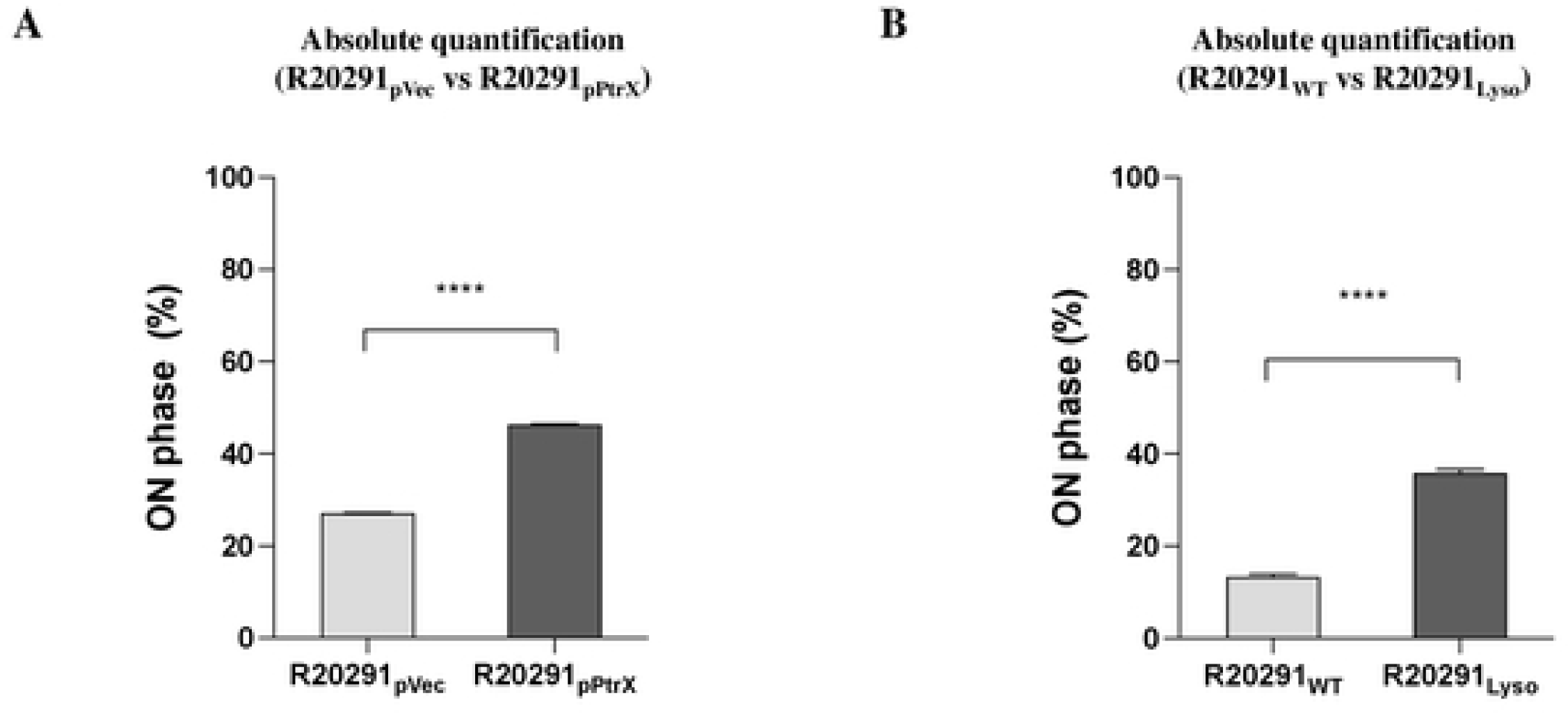
Absolute quantification of phase variation-associated states in R20291_Lyso_ and R20291_pPtrX_ strains. The absolute quantification of Cdi4 ON configuration was measured in the (A) R20291_Lyso_ and the (B) R20291_pPtrX_ strains, which were respectively compared with the R20291_WT_ and the vector control. All data were presented as the mean ± standard deviation values. At least three independent biological replicates of the experiment were conducted with similar results. Data from one representative independent biological replicate are shown. All statistical analyses were performed using the t-test analysis of variance test (\*\*\*\**p* ≤ 0.0001).

## Discussion

In this study, we successfully induced φNCKUH-21 by treating strain NCKUH-21 with mitomycin C. φNCKUH-21 infected the hypervirulent *C. difficile* strain R20291 and enhanced PaLoc gene expression and toxicity. The lysogenized R20291 exhibited a higher level of pathogenicity in swimming motility, spore germination, and biofilm formation than the WT strain. In addition, the lysogenized R20291 also caused a more severe infection in the CDI mouse model. We further investigated the mechanism involved in the φNCKUH-21-mediated regulation of PaLoc gene expression and pathogenicity in lysogenized R20291, and a putative transcriptional regulator, NCKUH-21_03890 (PtrX), plays a pivotal role in regulating the pathogenicity of *C. difficile*. Besides, PtrX modulated bacterial phase variation and *csrA* expression. We proposed mechanisms (Fig 9) underlying the φNCKUH21-mediated regulation of *C. difficile* pathogenicity. φNCKUH-21 produced PtrX, which directly or indirectly regulated the flagellar phase variation of *C. difficile* to increase the proportion of the Cdi4 phase ON, which could further modulate the expression of downstream *flgB* operon genes, to upregulate the expression of PaLoc genes. Notably, the upregulation of *csrA* expression by PtrX may enhance bacterial pathogenicity.

**Fig 9.**
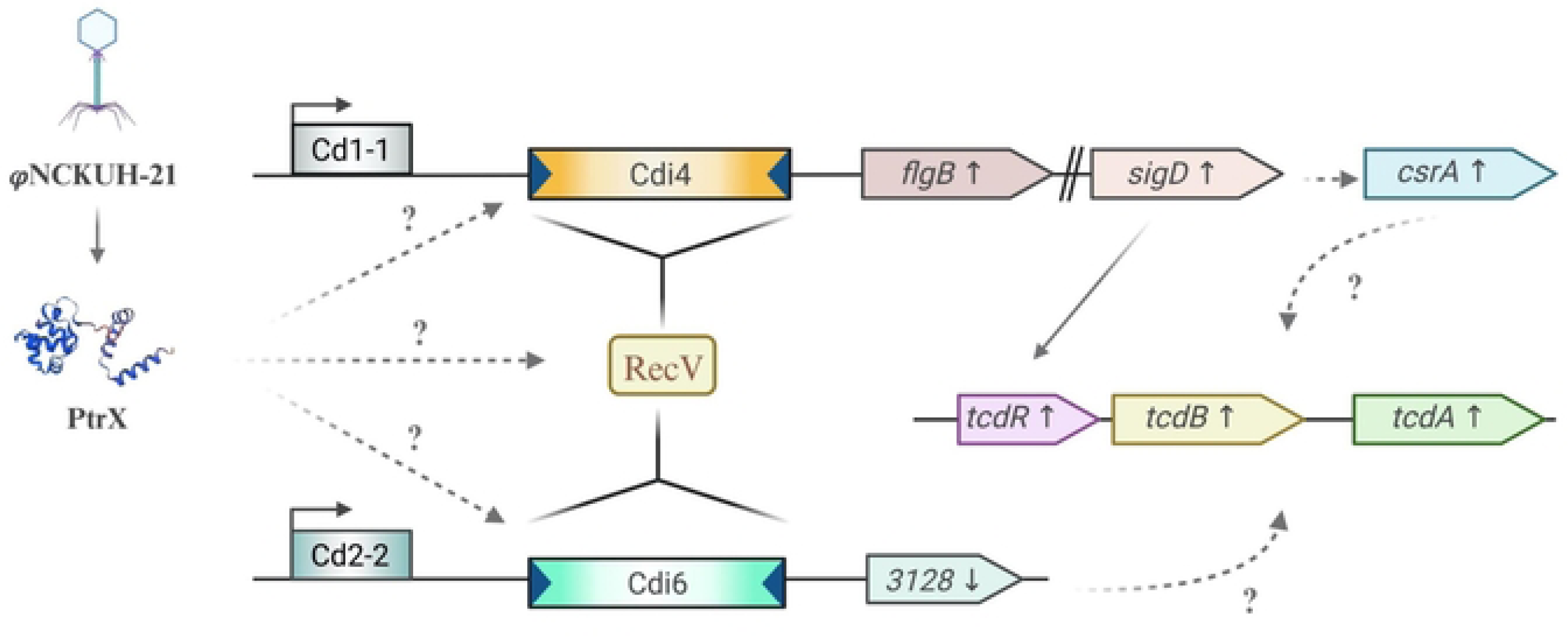
The proposed mechanism underlying the φNCKUH-21-mediated regulation of *C. difficile* pathogenicity. Schematic of a proposed mechanism underlying the φNCKUH21-mediated regulation of *C. difficile* pathogenicity (Created with BioRender.com). φNCKUH21 produced DNA-binding protein, PtrX, may interact with the invertible DNA regions or the tyrosine recombinase (RecV) to further influence the expression of *flgB*, *CDR20291_3128*, and *csrA* to enhance toxin production, bacterial motility, and biofilm formation. Cd1-1 and Cd2-2 indicate the c-di GMP riboswitch regions.

Infection with bacteriophages can induce the formation of lysogenized *C. difficile* cells; some prophages present in the bacterial host can be isolated (39, 45, 51). Prophages anchored in the *C. difficile* genome may regulate toxin production (2), and whether toxin production is increased or reduced depends on the anchoring prophage. φCD119 suppresses the expression of *tcdA* and *tcdB* in *C. difficile*. It reduces exotoxin production in the lysogenic strain of *C. difficile* through the RepR protein, which inhibits the transcription of *tcdR* (52, 53). φCD38-2 has been reported to stimulate toxin production in *C. difficile* RT027 strains. However, the mechanisms employed by φCD38-2 to enhance the toxin production of *C. difficile* remain unclear (2, 45).

The phage gene *ptrX* plays an important role in regulating the toxicity of *C. difficile* strains; therefore, we tried to use *ptrX* as the marker gene to search in the database, and the National Center for Biotechnology Information (NCBI) BLAST results revealed that approximately 500 *C. difficile* strains contain the prophage gene *ptrX.* We also detected three φNCKUH-21 genes (*NCKUH-21_03888, ptrX,* and *NCKUH-21_03903)* in *C. difficile* strains isolated from Chang Gung Memorial Hospital in northern Taiwan and Chi Mei Medical Center in southern Taiwan. These findings suggest that the presence of the prophage φNCKUH-21 in *C. difficile* strains is not an unusual event. However, more *C. difficile* strains should be studied to estimate the incidence of bacteria carrying φNCKUH-21.

The gene expression microarray analysis revealed that the expression levels of *tcdA* and *tcdB* were not upregulated in R20291_pPtrX_ transconjugant compared with in the vector control, it might cause by the harvesting of bacterial sample for gene expression microarray was at mid-log phase (OD_600_, approximately 0.4–0.6) which had no high toxins mRNA expression. Flagellin filament structural proteins (*fliC*), flagellar cap proteins (*fliD*), flagellar motor switch proteins (*fliG*, *fliM*, and *fliN*), and chemotaxis proteins (*motA* and *motB*) are necessary for flagellar motility (54-56).These genes are among DEGs list and upregulated in R20291_pPtrX_ transconjugant. This finding was consistent with the observed increase in the swimming motility of R20291_pPtrX_ transconjugant. Biofilm formation, another major contributor to *C. difficile* colonization, is associated with the expression of flagellar regulon genes, such as *fliC* in R20291 (57). Several flagellar proteins are detected in high concentration in aggregate biofilm, such as FliE, FliF, FliG, FliM, FlgE, and FlgG (58). These genes are among the DEGs upregulated in R20291_pPtrX_ transconjugant as compared to the vector control. This finding corroborates enhanced biofilm formation in both the lysogenized and transconjugant strains. The overexpression of *csrA* could increase toxin production has been reported in *C. difficile* 630Δerm (27), but no study in R20291 has been conducted. Therefore, whether the upregulation of toxin genes contributed by a higher expression level of *csrA* in R20291_pPtrX_ remains unclear. Bacterial phase variation is essential for regulating the expression toxin-related and flagellar genes, and the ratio of Cdi4 phase ON to phase OFF configuration must be regulated to modulate toxin production (17). Taken together, the observations indicate that PtrX enhances the toxicity and pathogenicity of R20291 by upregulating the expression of flagellar genes.

Recombinase-regulated flagellar phase variation mediates the link between *C. difficile* flagella and toxin production and pathogenesis (54, 56, 59). It has been reported that a lysogenic strain of *C. difficile* carrying φCD38-2 exhibited increased *cwpV* expression (45). However, we found no significant difference in the expression levels of *cwpV* between R20291_pPtrX_ transconjugant and the vector control. The difference between the two study results may be due to the different experiment designs, the comparison in this study reveals the effects of a single phage gene. The expression of *flgB* and its downstream gene *sigD* was significantly upregulated in R20291_pPtrX_ transconjugant. Furthermore, the expression of Cdi6 was significantly downregulated in R20291_pPtrX_; the inverted orientation may represent the phase ON (15, 17). Our findings indicate that PtrX regulates the expression of *csrA* and the genes responsible for the flagellar phase variation of *C. difficile*, thus enhancing the pathogenicity of lysogenized R20291 carrying the prophage φNCKUH-21. Although the absolute quantification results revealed that PtrX significantly increased the percentage of phase ON in lysogenized R20291 cells, the precise mechanisms involved in the regulation of phase variation remain to be elucidated. RecV, a tyrosine recombinase that regulates flagellar switch inversion in *C. difficile*, can modulate the phase ON/OFF configuration ratio (15, 18). However, further studies are needed to validate the association between PtrX and RecV. Notably, PtrX, a putative CI repressor that can bind to DNA, may directly interact with the promoter regulated by flagellar phase variation. We are currently investigating whether PtrX can directly bind to a flagellar switch–specific DNA sequence. The results of the gene expression microarray analysis performed in the present study indicated that the expression of *csrA* was upregulated in the presence of PtrX. However, the association between the expression of *csrA* and the upregulation of the genes related to flagellar phase variation remains to be identified. In the future, we may overexpress CsrA in R20291 to monitor the changes in the phase ON/OFF configuration ratio to confirm the association. In addition, the DNA sequence of three φNCKUH-21 genes tested in this study is 100% identical to the corresponding genes of φCD38-2. Therefore, the results in this study may also apply to φCD38-2. Overall, the study provides insights into how the φNCKUH-21 bacteriophage interacts with *C. difficile* R20291 and how specific genes within the phage genome can influence the expression of pathogenicity-related genes, bacterial motility, biofilm formation, and infection severity.

## Materials and Methods

### Bacterial strains and culture conditions

All strains of bacteria used in this study are listed in S1 Table. The *C. difficile* strains were cultured anaerobically (10% H_2_, 10% CO_2_, and 80% N_2_) at 37℃ on BHI agar or in BHI broth (Thermo Fisher Scientific, Waltham, MA, USA) supplemented with 0.05% L-cysteine and yeast extract and tryptone–yeast extract (TY) medium (3% tryptone, 2% yeast extract, and 0.1% thioglycolate). Anaerobic bacteria were maintained in an anaerobic workstation (DG250; Don Whitley Scientific Ltd., West Yorkshire, UK). All *E. coli* strains used in this study were cultured aerobically at 37℃ on Luria–Bertani (LB) agar or in LB broth. All media were supplemented with 15 μg/mL thiamphenicol, 300 μg/mL cycloserine, 12.5 μg/mL chloramphenicol, 100 μg/mL ampicillin, and 50 μg/mL kanamycin, as necessary.

### Prophage induction

NCKUH-21 cells were inoculated in 7 mL of BHIS broth and anaerobically cultured at 37℃ till the stationary phase of growth was attained (OD measured at 600 nm [OD_600_], approximately 0.8). The overnight culture of NCKUH-21 cells was diluted 100-fold in a fresh BHIS medium. Mitomycin C (MDBio Inc., Taiwan) (46) was added to the final concentration from 2 to 5 μg/mL when the OD_600_ value reached approximately 0.1. The mitomycin C-containing culture was anaerobically incubated at 37°C; the OD_600_ value was measured once per hour. A marked decrease in cell density after 3–5 h indicated phage release. Then, the culture was centrifuged at 2,200 ×*g* for 15 min at 25℃. The supernatant was filtered through a 0.45-μm filter and stored at 4°C until further analysis.

### Bacteriophage infection

Overnight anaerobic culture (BHIS broth, 37°C) of R20291 cells was diluted 100-fold in fresh BHIS medium and grown till the late exponential to early stationary phase (OD_600_, approximately 0.8). The cultures of R20291 were mixed with φNCKUH21 solution at the ratios of 1:1, 1:2, and 1:3, respectively. To enhance infection efficiency, CaCl_2_ and MgCl_2_ were added (final concentration, 10 mM) to each bacteria–phage mixture (60). The mixtures were anaerobically incubated at 37℃ for 2 h and streaked on BHIS agar. Eight colonies were selected to perform PCR with the primers of *gp39* (S2 Table) to check the presence of φNCKUH-21 in colonies.

### Genomic DNA characteristic of φNCKUH-21

The complete genomic coding sequence (CDS) of φNCKUH-21 was assembled with BLAST Ring Image Generator software (BRIG) (61). Functional proteins were annotated using NCBI BLASTp databases to compare the coding region translated amino acid sequences of φNCKUH-21.

### Transmission electron microscopy (TEM)

The TEM sample was prepared as previously described (62). In brief, 5 μL of filtered φNCKUH-21-induction supernatant was dropped on the TEM grid (Ted Pella Inc., USA) and incubated for 1 min at 25°C. The remaining phage solution was removed, and the sample was then fixed for 30 sec using 10 μL 2% paraformaldehyde (Sigma Aldrich, USA). Following that, the grid was washed three times with 15 μL ddH_2_O and negatively stained with 2% OsO4 (Sigma Aldrich, USA) and 2% uranyl acetate (Sigma Aldrich, USA). After performing wash and air-dry, the TEM images of φNCKUH-21 were acquired using Zeiss EM 912 TEM (Zeiss, Germany)

### Biofilm formation assay

The WT, lysogenized, and transconjugant R20291 strains of *C. difficile* were cultured overnight in 6 mL BHIS broth containing 0.1 M glucose. The overnight culture was diluted 100-fold in fresh medium in 24-well plates (GeneDirex, Miaoli, Taiwan) and grown till the OD_600_ value reached approximately 1.0. Then, the plates were anaerobically incubated at 37°C for 5 days; the plates were wrapped with parafilm to prevent the loss of medium. After incubation, the corresponding supernatants were carefully collected for the quantification of the biofilm mass. The plates were dried at 25℃ for 30–60 min, followed by the addition (to each well) of 500 μL of 2% crystal violet and incubation at 25℃ for 30 min. After the removal of the dye, 500 μL of methanol was added to each well and incubated on an orbital shaker (model 022057B; Major Science, Taoyuan, Taiwan) for 30 min. All methanol-extracted dye contents were transferred to 1.5-mL Eppendorf tubes and centrifuged at 17,900 ×*g* for 1 min. The supernatant (200 μL) was collected from each sample and transferred to 96-well ELISA plates for the measurement of the absorbance at 595 nm by using the iMark^TM^ Microplate Absorbance Reader (Bio-Rad, Alfred Nobel Drive. Hercules, CA, USA).

### Bacterial RNA extraction

*C. difficile* strains were anaerobically cultured overnight in TY broth. The overnight cultures were diluted 100-fold in fresh medium and grown till the mid-log phase was attained (OD_600_, approximately 0.4–0.6). The RNAprotect Bacteria Reagent (500 μL; Qiagen, Hilden, Germany) was added to each bacterial culture (4 mL) and incubated at 25℃ for 5 min. Then, the samples were centrifuged at 2,200 ×*g* for 15 min at 4℃. The corresponding pellets were stored at −80℃. Total RNA was extracted using the RNeasy Mini Kit (Qiagen) according to the manufacturer’s instructions. First-strand cDNA was synthesized from total RNA (3 μg) by using the SuperScript III First-Strand Synthesis System (Invitrogen, Van Allen Way, Carlsbad, CA, USA) and random primers (Invitrogen) according to the manufacturer’s instructions. The obtained cDNA samples were diluted to a final concentration of 20.0 ng/μL.

### Bacterial gene expression microarray

Gene expression microarray was performed by Agilent Technologies (Santa Clara, CA, USA). In brief, overnight anaerobic cultures (in BHIS) of *C. difficile* transconjugants were diluted 100-fold in fresh BHIS medium and grown till the OD_600_ value reached approximately 0.6. Then, each cell culture was centrifuged at 2,200 ×*g* for 15 min at 4℃. The pellet was then resuspended in 500 μL of the RNAprotect Bacteria Reagent and incubated at 25℃ for 5 min. The suspension was centrifuged at 2,200 ×*g* for 15 min at 4℃. The pellet was stored at −80℃. From the *C. difficile* transconjugants, total RNA was extracted using the RNeasy Mini Kit following the manufacturer’s instructions. Total RNA (0.2 μg) was amplified using the Low Input Quick Amp Labeling Kit (Agilent Technologies) and labeled with Cy3 (CyDye; Agilent Technologies) during in vitro transcription. Cy3-labeled cRNA (0.6 μg) was incubated in a fragmentation buffer at 60°C for 30 min and was thus fragmented into 50–100-bp nucleotides. Next, the labeled cRNA fragments were pooled and hybridized to an Agilent SurePrint Microarray (Agilent Technologies) at 65°C for 17 h. After washing and drying (nitrogen gun blowing), the microarrays were scanned at 535 nm (for Cy3) by using a microarray scanner (Agilent Technologies). The obtained images were analyzed using Agilent Feature Extraction Software (version 10.7.3.1; Agilent Technologies), which is an image analysis and normalization software program used to quantify the signal and background intensity of each feature. Raw signal data were normalized through quantile normalization to identify differentially expressed genes. To assess gene functions, enrichment analyses were performed (at Welgene Biotech) for differentially expressed genes (for most model organisms). The cluster Profiler tool was used for gene ontology and Kyoto Encyclopedia of Genes and Genomes pathway analyses. For data analysis, we selected the genes whose median expression was upregulated or downregulated (at least by 2-fold) in the R20291_p3890_ transconjugant compared with the expression levels in the R20291_pVec_ transconjugant.

### Bacterial gene expression analysis with RT-qPCR

The expression levels of PaLoc genes were measured through RT-qPCR. The reaction mixture comprised the cDNA template (20.0 ng); specific primer sets for *tcdA*, *tcdB*, *tcdC*, *tcdR* and *csrA* (200 nM); and 2× GoTaq qPCR Master Mix (Promega, Madison, WI, USA). RT-qPCR was performed using the MyGo Pro real-time PCR instrument (IT-IS Life Science Ltd., Cork, Ireland). The cycling conditions were as follows: initial denaturation at 95°C for 10 min, followed by 40 cycles of denaturation at 95°C for 30 s and annealing and extending at 60°C for 1 min. The obtained data were analyzed using the 2^–ΔΔCt^ method (63). The expression levels of the test genes were normalized to the expression level of the 16S rRNA gene (64). S2 Table lists the primers used in this study. All data were representative of at least two independent experiments and each experiment was performed in quadruple.

### Bacterial genomic DNA extraction and absolute quantification in qPCR

Overnight anaerobic cultures (in BHIS) of *C. difficile* WT R20291, lysogenized R20291 and R20291 transconjugants were diluted 100-fold in fresh BHIS medium and grown till the OD_600_ value reached approximately 0.6. Then, each cell culture was centrifuged at 2,200 ×*g* for 15 min at 4℃ to harvest bacterial pellet. Genomic DNA was extracted following the Presto^TM^ Mini gDNA Bacteria Kit Quick Instruction Manual (Geneaid Biotech Ltd, New Taipei City, Taiwan). In brief, bacterial pellet was mixed with lysozyme-Gram^+^ Buffer and incubated at 37℃ for 30 mins. Add 20 µL of Proteinase K (10 mg/mL) then mix by vortex and incubate at 60℃ for 10 mins. Next, 200 µL of GB buffer was added to lysis bacterial pellet and GD column was used to perform the DNA binding and washing steps. The purified genomic DNA was then eluted with nuclease-free H_2_O.

Specific primer sets, forward 5’AAACTATGTCAGCCAGTTGCC3’ and reverse 5’AGGCATAGCATCATTTAGTGTTTCTTC3’ (18), were used to amplify the *flgB* upstream phase variation region and CloneJET PCR Cloning Kit (Thermo Fisher Scientific Inc.) was used to create a vector that carrying phase ON (pJET-ON) or OFF (pJET-OFF) fragments as the template to perform the standard curve of absolute quantification. The copy number of phase variation was measured through qPCR. The reaction mixture comprised the gDNA template (20.0 ng); specific primer sets for phase ON/OFF (30 µM) (18); and 2× GoTaq qPCR Master Mix (Promega). qPCR was performed using the MyGo Pro real-time PCR instrument (IT-IS Life Science Ltd.). The cycling conditions were as follows: initial denaturation at 95°C for 10 min, followed by 40 cycles of denaturation at 95°C for 45 s and annealing and extending at 60°C for 1 min. The obtained data were analyzed using the absolute quantification method with standard curves (S5 Fig). All data were representative of at least two independent experiments and each experiment was performed in quadruple.

### Plasmid construction and transformation

The genes *NCKUH-21_03888*, *NCKUH-21_03890*, and *NCKUH-21_03903* (including the endogenous promoter) were amplified through PCR performed using the genomic DNA of NCKUH-21 cells and specific primers (3888-XbaI-F/3888-EcoRI-R, 3890-XbaI-F/3890-EcoRI-R, and 3903-XbaI-F/3903-EcoRI-R, respectively). The PCR products were digested with *Eco*RI and *Xba*I and then cloned into the pMTL-84151 vector to generate plasmids, which were further confirmed through DNA sequencing. Next, the plasmids were transformed into the *E. coli* strain CA434 (plasmid donor). Through conjugation, the plasmids were transferred from the donor bacteria to strain R20291.

### Competent cell preparation

Overnight culture (LB, 37°C) of a single *E. coli* colony was diluted 50-fold in fresh LB broth and grown till the OD_600_ value reached 0.5. The bacteria were incubated on ice for 30 min and centrifuged at 4,000 ×*g* for 25 min at 4°C. The pellet was resuspended in 5 mL of transformation and storage solution (TSS; 10% polyethylene glycol [molecular weight, 3,350], 5% dimethyl sulfoxide, 10 mM MgCl_2_, and 10 mM MgSO_4_ in LB broth). The prepared competent cells were stored at −80℃.

### Transformation into *E. coli*

The R20291_p3888_, R20291_p3890_ and R20291_p3903_ plasmids were suspended in solution containing 20 μL of 5× KCM buffer (0.5 M KCl, 0.15 M CaCl_2_, and 0.25 M MgCl_2_) and 80 μL of sterile double-distilled water (ddH_2_O). Subsequently, 100 μL of competent cells was added to this suspension and incubated on ice for 20 min. Following incubation, the mixture was subjected to heat shock (water bath at 42℃) for 1 min and then quickly placed on ice for 1 min. For bacterial recovery, 1 mL of precooled LB broth was added to the mixture and incubated at 37℃ for 1 h. Next, the culture was centrifugated at 10,000 ×*g* for 1 min at 25℃; the pellet was collected. The pellet was resuspended with 100 μL fresh LB broth and inoculated on LB agar containing an appropriate antibiotic for transformant selection.

### Conjugation of *C. difficile* with *E. coli*

Overnight culture of the CA434 transconjugant donor was centrifuged at 4,000 ×*g* for 2 min at 25℃. The collected pellet was gently washed three times with 1 mL of anaerobic BHI broth. Overnight culture of R20291 cells (plasmid acceptor) was heated at 50℃ for 5 min and then centrifuged at 2,200 ×*g* for 2 min at 25℃. The collected pellet was resuspended in 1 mL anaerobic BHI broth. The suspension was anaerobically incubated for 4 h and then added in a dropwise manner onto BHI agar plates (20 μL × 10 spots). The bacterial spots were anaerobically incubated for 4 h. Next, the spots were suspended in 800 μL of anaerobic BHIS broth and then spread on BHIS agar containing 250 μg/mL cycloserine and 15 μg/mL thiamphenicol for conjugant selection. All colonies were confirmed through PCR performed using specific primers for *C. difficile* (tpi-F/tpi-R) and NCKUH-21 (3890-NdeI-F/3890-XhoI-R).

### ELISA for toxins A and B

To detect *C. difficile* toxins, all strains were anaerobically cultured in TY broth overnight at 37℃. The overnight cultures were diluted 100-fold in 10 mL of anaerobic fresh TY broth and grown for 30 h. Then, each culture was centrifuged at 2,200 ×*g* for 15 min. The supernatant was collected and filtered through a 0.45-μm filter. Toxin production was assessed using an enzyme immune assay kit (Premier Toxins A&B; Meridian Bioscience, Cincinnati, OH, USA) following the manufacturer’s instructions. In brief, each filtered bacterial supernatant was five folds diluted and mixed with 50μL of Enzyme Conjugate solution for 20 mins at 37°C. After washing 4 to 6 times, 100μL of Substrate I was dropped to all wells and incubated for 10 mins at 25°C. After the addition of the stop solution (provided in the kit), the reaction mixture was incubated for 10 min, which was followed by the measurement of absorbance at 450 nm by using the iMark^TM^ Microplate Absorbance Reader. This experiment was representative of three independent experiments and each experiment was performed in triplicate.

### Cell culture and cytotoxicity assay

Caco-2 cells were cultured at 37°C in Dulbecco’s Modified Eagle Medium/Nutrient Mixture F-12 (GenedireX) supplemented with 1× antibiotic solution (10,000 units/mL Penicillin, 10,000 μg/mL Streptomycin, 25 μg/mL Amphotericin B; Thermo Fisher Scientific Inc.). The cells were detached using trypsin–ethylenediaminetetraacetate, stained with trypan blue, and counted using a cell counter. Caco-2 cells were seeded (density of 3 × 10^4^ cells/well) onto 96-well plates (SPL life sciences, Pocheon, South Korea) and incubated at 37°C for 16 h under 5% CO_2_. For cytotoxicity assay, all *C. difficile* strains were anaerobically cultured in TY broth overnight at 37°C. The overnight cultures were diluted 100-fold in 10 mL fresh TY broth and grown till the OD_600_ value reached 1.2. Next, each culture was centrifuged at 2,200 ×*g* for 15 min, and the supernatant was collected. To the supernatant, 10% filtered glycerol was added. For the cell viability assay, Caco-2 cells were seeded (3 × 10^4^ cells/well) onto a 96-well tissue culture plates and incubated overnight at 37°C under 5% CO_2_. Caco-2 cells were coincubated for 18 h in *C. difficile* condition medium diluted in anaerobic TY broth. Then, Caco-2 cells were detached using trypsin–ethylenediaminetetraacetate and then stained with trypan blue (Logos Biosystems, Anyang, South Korea). Finally, living cells were counted using the LUNA-FL Dual Fluorescence Cell Counter (Logos Biosystems). This experiment was representative of three independent experiments and each experiment was performed in triplicate.

### Experimental CDI animal model and sample collection

Inbred male C57BL/6 mice (age, 7 weeks) were purchased from the Laboratory Animal Center of NCKU. All mice were maintained and handled in accordance with the guidelines of the Institutional Animal Care and Use Committee (IACUC) of NCKU. Moreover, all animal studies were performed following the protocol approved by the IACUC of NCKU (approval number: NCKU-IACUC-107-092) and the NCKU Biosafety and Radiation Safety Management Division. To establish a mouse model of CDI, an antibiotic cocktail (Sigma-Aldrich, St. Louis, USA) comprising vancomycin (0.4 mg/mL), metronidazole (0.215 mg/mL), kanamycin (0.4 mg/mL), gentamycin (0.035 mg/mL), and colistin (850 U/mL) was administered to 8-week-old mice through drinking water 4 days before the challenge with different strains of *C. difficile*. A day before infection, vancomycin and metronidazole were eliminated from the cocktail to avoid the disruption of bacterial colonization. Mice were intraperitoneally injected with 100μL of 1 mg/mL clindamycin and were orally inoculated with (1) sterile 0.9% normal saline (control; Mock group, 11 mice), (2) PBS containing 10^7^ colony-forming units (CFUs) of WT strain R20291 (R20291_WT_ group, 11 mice), or (3) PBS containing 10^7^ CFU of lysogenized R20291 (R20291_Lyso_ group, 14 mice). All mice were euthanized through CO_2_ asphyxia 3 days after infection. The organs were harvested and stored at -80°C for downstream analyses. Resected colon tissue samples were fixed in 10% formaldehyde (BS Chemical, Kretinga, Lithuania) buffered with PBS and then embedded in paraffin. Tissue sections were stained with hematoxylin and eosin (H&E). Animal experiments were representative of two independent experiments and each experiment was performed in at least triplicate.

### Histopathology and colitis severity scoring

The paraffin-embedded colonic tissue samples were sectioned to 5 μm and performed H&E staining by SIDSCO Biomedical Co., Ltd (Kaohsiung, Taiwan). The stained sectioned tissue samples were imaged through microscopy for determining tissue integrity. For pathological scoring, six high-power fields (HPFs) per tissue sample were examined and scored. The average numbers of neutrophils in the HPFs were quantified. The severity of colitis was scored. The score ranged between 0 and 3 for each of the following parameters: polymorphonuclear infiltrate, mononuclear infiltrate, edema, erosion and ulceration, crypt abscess, crypt destruction, and inflammation distribution (mucosa = 1; mucosa and submucosa = 2; and transmural inflammation = 3) (65-67). Various inflammation markers, including neutrophil infiltration, epithelial damage, and irregularly shaped colon mucosa, were analyzed.

### Spore purification

Overnight cultures of *C. difficile* were diluted 100-fold in fresh BHIS broth and grown till the OD_600_ value reached 0.6–0.8. Then, the cells were plated on an agar medium containing 70% smooth muscle cell medium and 30% BHIS medium (68). Following anaerobic incubation at 37℃ for 7–9 days, bacterial spores were harvested in ice-cold sterile ddH_2_O and stored overnight at 4°C. The spores were centrifuged at 2,330 ×*g* for 16 min at 4°C and washed five times with ice-cold sterile ddH_2_O to separate spores from the debris and vegetative cells. Next, the spores were purified using the universal density gradient medium Nycodenz (48%; Axis Shield, Oslo, Norway) and centrifuged at 2,500 ×*g* for 10 min. The pellet containing the spores was collected and washed five times with ice-cold sterile ddH_2_O. This was followed by centrifugation at 2,500 ×*g* for 7 min at 25°C. The pellet was resuspended in PBS and stored at 4°C for subsequent experiments.

### Sporulation assay

Overnight cultures of *C. difficile* were diluted 100-fold in fresh BHIS broth and anaerobically incubated at 37℃ incubated for 5 days. For the counts of total vegetative cells, *C. difficile* cultures were 10-fold serially diluted and grown on BHIS agar supplemented with 0.1% TA. For the counts of purified spore, the cultures were treated with 30 min incubation at 65 °C before serial dilution. The colonies were counted after 24 h incubation at 37 °C. The percentage of sporulation was calculated by spore CFUs/ total cell CFUs (69, 70)

### Spore germination assay

The percentage of completely germinated *C. difficile* spores was evaluated by estimating the rate of colony formation. The spore suspensions were quantified, serially diluted, and plated onto BHIS agar supplemented with 0.1% taurocholate and anaerobically incubated at 37℃ for 24 h. The concentration of mature spores in the suspensions was directly measured on a hemocytometer (Improved Neubauer) under a phase-contrast microscope. Germination efficiency was defined as the percentage of mature spores that formed colonies and was calculated by dividing the number of CFU on agar plates by the number of spores quantified through microscopy (71, 72).

### Bacterial tube and soft agar motility assays

Motility assays were performed using the stab culture technique (73, 74) and soft agar swimming assay (18) following the previously described approaches. Overnight cultures from various *C. difficile* strains (R20291_WT_, R20291_Lyso_, R20291_pVec_ and R20291_pPtrX_) in BHIS broth were diluted 100-fold in fresh broth and grown till the OD600 value reached 1.0. For tube motility, refreshed cultures were vertically stab-inoculated in BHIS-0.175% agar and anaerobically incubated at 37°C for 24 h. The areas of bacterial growth were measured using ImageJ (version 8) and the whole tube area was used as 100% for normalizing bacterial motility. For soft agar motility assays, 5 μL of refreshed cultures were spotted onto BHIS-0.3% agar and anaerobically incubated at 37°C. The diameters of growth were measured at 24-hour intervals for 3 days.

### Bioinformatics analysis

The DEGs list obtained from microarray was analyzed using STRING version 12.0 (https://string-db.org/) to perform protein-protein interaction networks functional enrichment analysis and gene ontology (GO) annotation enrichment analysis.

## Acknowledgments

This work was supported by the National Science and Technology Council (Grants: 107-2320-B-006-043, 108-2320-B-006-036, and 111-2314-B-006-007; awarded to JWC). The funding agency played no role in study design, data collection or analysis, manuscript preparation, or decision to publish. We thank the Laboratory Animal Center, College of Medicine at National Cheng Kung University (NCKU), Taiwan, accredited by AAALAC International and Taiwan Animal Consortium. We appreciate the support of materials from Dr. Chang-Shi Chen, Department of Biochemistry and Molecular Biology, National Cheng Kung University. This manuscript was edited by Wallace Academic Editing.

## Data availability statement

All data performed in this study are available in the article and its supplementary materials. The gene expression microarray data has been deposited in Gene Expression Omnibus (GEO) with accession number GSE219066.

